# MASCOT: Parameter and state inference under the marginal structured coalescent approximation

**DOI:** 10.1101/188516

**Authors:** Nicola F. Müller, David A. Rasmussen, Tanja Stadler

## Abstract

**Motivation:** The structured coalescent is widely applied to study demography within and migration between sub-populations from genetic sequence data. Current methods are either exact but too computationally inefficient to analyse large datasets with many states, or make strong approximations leading to severe biases in inference. We recently introduced an approximation based on weaker assumptions to the structured coalescent enabling the analysis of larger datasets with many different states. We showed that our approximation provides unbiased migration rate and population size estimates across a wide parameter range.

**Results:** We here extend this approach by providing a new algorithm to calculate the probability of the state of internal nodes that includes the information from the full phylogenetic tree. We show that this algorithm is able to increase the probability attributed to the true node states. Furthermore we use improved integration techniques, such that our method is now able to analyse larger datasets, including a H3N2 dataset with 433 sequences sampled from 5 different locations.

**Availability:** The here presented methods are combined into the BEAST2 package MASCOT, the Marginal Approximation of the Structured COalescenT. This package can be downloaded via the BEAUti package manager. The source code is available at https://github.com/nicfel/Mascot.git.

## 1 Introduction

Phylogenies obtained from a multiple sequence alignment, contain information regarding the history of a population and can be used to quantify demographic parameters. This has been widely done to study the spread of pathogens (Pybus et al., 2001; Russell et al., 2008), the speciation dynamics of extant species or the migration pattern of humans to name but a few. Forwards in time birth-death and backwards in time coalescent based models allow us to elucidate population dynamics from trees by calculating the probability of a phylogeny *T* given a set of demographic parameters Θ. To do so they classically rely on the assumption of well mixed populations, meaning that the probability of any two pairs of lineages to share a common ancestor is the same. In most empirical applications this assumption of well mixed populations is however violated.

To address this model violation, so-called structured methods have been developed that consider birth-death processes in heterogeneous populations (Stadler and Bonhoeffer, 2013). In the backward-in-time coalescent framework, the structured coalescent (Takahata, 1988; Hudson, 1990; Notohara, 1990) describes a coalescent process in sub-populations between which individuals can migrate. Such coalescent methods however typically require the state (or location) of any ancestral lineage in the phylogeny at any time to be inferred (Beerli and Felsenstein, 2001; Ewing et al., 2004; Vaughan et al., 2014). Inferring lineage states is computationally expensive, as it normally requires MCMC based sampling, and limits the complexity of scenarios that can be analysed. As the number of different states is increased, convergence of the MCMC chains becomes a severe issue (De Maio et al., 2015). This essentially limits the number of different states that can be accounted for to three or four.

We addressed this limitation recently by introducing a new approximation of the structured coalescent that avoids this MCMC sampling of lineage states by integrating over all possible migration histories using a set of ODEs (Müller et al., 2017). In contrast to previous approximations that treat the movement of one lineage completely independently of all other lineages (Volz, 2012; De Maio et al., 2015), we explicitly include information about the location of other lineages and their probability of coalescing when modelling the movement of a lineage. We showed that this approximation is able to infer coalescent and migration rates well in various scenarios. However, this approach currently lacks the possibility to estimate the ancestral state of any internal nodes except the root.

Here, we introduce a new algorithm to calculate the probability of internal nodes being in any state that incorporates information from the entire tree using a forwards/backwards approach (Pearl, 1982). We additionally make improvements of the current BEAST2 (Bouckaert et al., 2014) implementation of Müller et al. (2017) in terms of calculation speed, allowing larger datasets to be analysed. We then show first on simulated datasets how this new implementation performs in inferring migration rates and effective population sizes in high dimensional parameter space. Next, we show how our new algorithm can dramatically improve ancestral state inference. We then apply our new approach to a geographically distributed samples of human Influenza A/H3N2 virus to demonstrate its applicability to large datasets.

## 2 Materials and Methods

### 2.1 The approximate structured coalescent

In (Müller et al., 2017), we introduced a new approximation to the structured coalescent that integrates over every possible migration history and avoids the sampling of lineage states. This is done by calculating the marginal probability of a lineage *i* being in any of *m* possible states, jointly with the probability of having observed the coalescent history *T* from the present backwards in time until time point *t* in the tree, with time 0 being the time of the most recent sample with time increasing into the past. To do so, we need to make the following approximation:

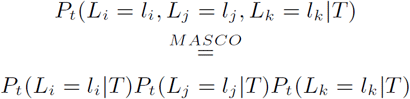

In other words, we assume that lineages *i, j* and *k* and their states *l_i_, l_j_* and *l*_*k*_ are pairwise independent.

### 2.2 The probability of a lineage being in a state

As described in (Müller et al., 2017), we seek to calculate the probability of every lineage being in any state jointly with the probability of having observed the coalescent history *T* up to time *t*. We previously denoted this probability as *P*_*t*_(*L*_*i*_ = *l_i_, T*). Calculating these terms over time for increasing t leads to ever smaller values, eventually causing numerical issues. To avoid this, we can calculate 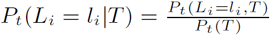 instead. The expression for 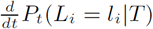 can be directly derived from 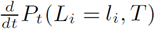 (see Supplement) and can be written as:

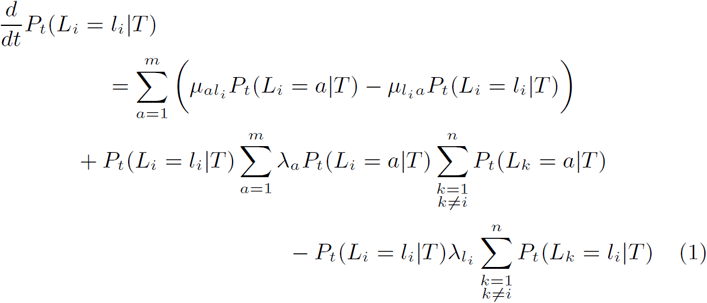

with 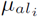 denoting the backwards in time rate at which lineages migrate from state *a* to state *l*_*i*_ and *λ*_*a*_ denoting the rate of coalescence in state *a*. To calculate *P*_*t*_(*T*), i.e. the probability of having observed the coalescent history *T* up to time *t*, the following differential equation has to be solved (see supplement for derivation):

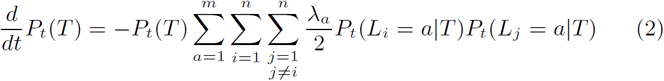

At coalescent events, we update *P*_*t*_(*T*) by multiplication with the probability of the coalescent event:

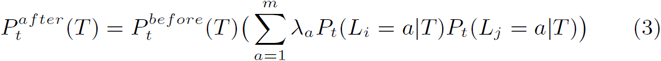

Integrating these equations from the present to the root of a phylogeny, allows to calculate *P*_*root*_(*T*), that is the probability density of a phylogeny

*T* under the MASCO approximations of the structured coalescent.

### 2.3 The probability of a node being in a state given the whole phylogeny

#### 2.3.1 Backwards calculation of node states conditional on the sub-trees

Integrating equation 1 allows to calculate the probability of each lineage being in any state given the coalescent history between the lineage and the present. However, for applications, it is much more interesting to calculate the probability of each lineage at time t being in any state given the whole phylogenetic tree between the time of root and the present. At coalescent events between lineage *i* and *j* at time t, the probability of the parent lineage *p* at time t being in state *a* can be calculated as follows:

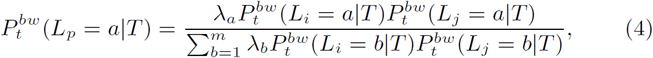

with 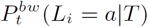 denoting the probability of the daughter lineage *i* being in state *a* just before the coalescent event at time t calculated in the backwards step using equation 1. Since 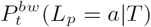 denotes the probability of the parent node of lineages *i* and *j* calculated in the backwards step at time t, it includes only information up to the time of coalescence and does not include information from the full phylogeny. We introduce the label “bw” to differentiate from the forward probabilities introduced below. To additionally incorporate information from the phylogeny between the time of the root and time t of the coalescent event, one has to deploy a backwards/forwards approach that is related to Pearl (1982).

For convenience, we now change to vector notation. We define 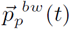 as the vector for the parent lineage *p* with entries 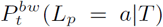 in position *a* that only includes information from time 0 up to time *t*. 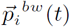 is the vector with entries 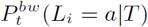.

#### 2.3.2 Calculation of transition probabilities

Going through the tree backwards in time, we also seek to calculate the probability that, given lineage *i* was in state *a* at the last coalescent event *lc* were lineage *i* was the parent and given the tree *T* between 0 and time *t*, the lineage i is ending up in state *b* at the time of the next coalescent event *nc* involving lineage *i*. We denote this probability as *P*_*t*=*nc*_(*L*_*i*_ = *b*|*L*_*i*(*t*=*lc*)_ = *a*). The Matrix *M*_*i*_ with entries *P*_*t*=*nc*_(*L*_*i*_ = *b*| *L*_*i*(*t*=*lc*)_ = *a*) in positions (*a, b*) now denotes the matrix for which the following equation holds:

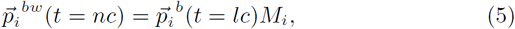

with 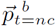 being the vector with the state probabilities of lineage *i* just before its next coalescent event *nc*. To calculate the entries of the matrix *M*_*i*_, we solve the following differential equation between the two coalescent events *lc* and *nc* involving lineage *i*:

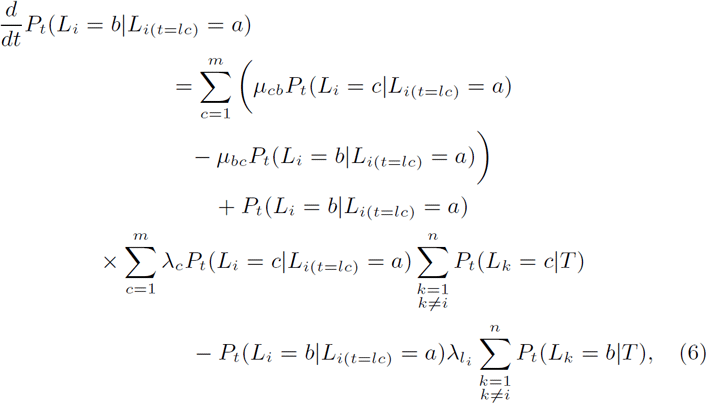

The entries in positions (*a, b*) of the Matrix *M*_*i*_ are then the solution of the above differential equation at time *nc* with initial values 1 if *b* = *a* and 0 otherwise. In other words, equation 6 describes solving equation 1 with lineage i starting in a (rather than in any state as in equation 1), assuming that all other lineage ≠ *i* evolve according to equation 1.

**Figure 1:**
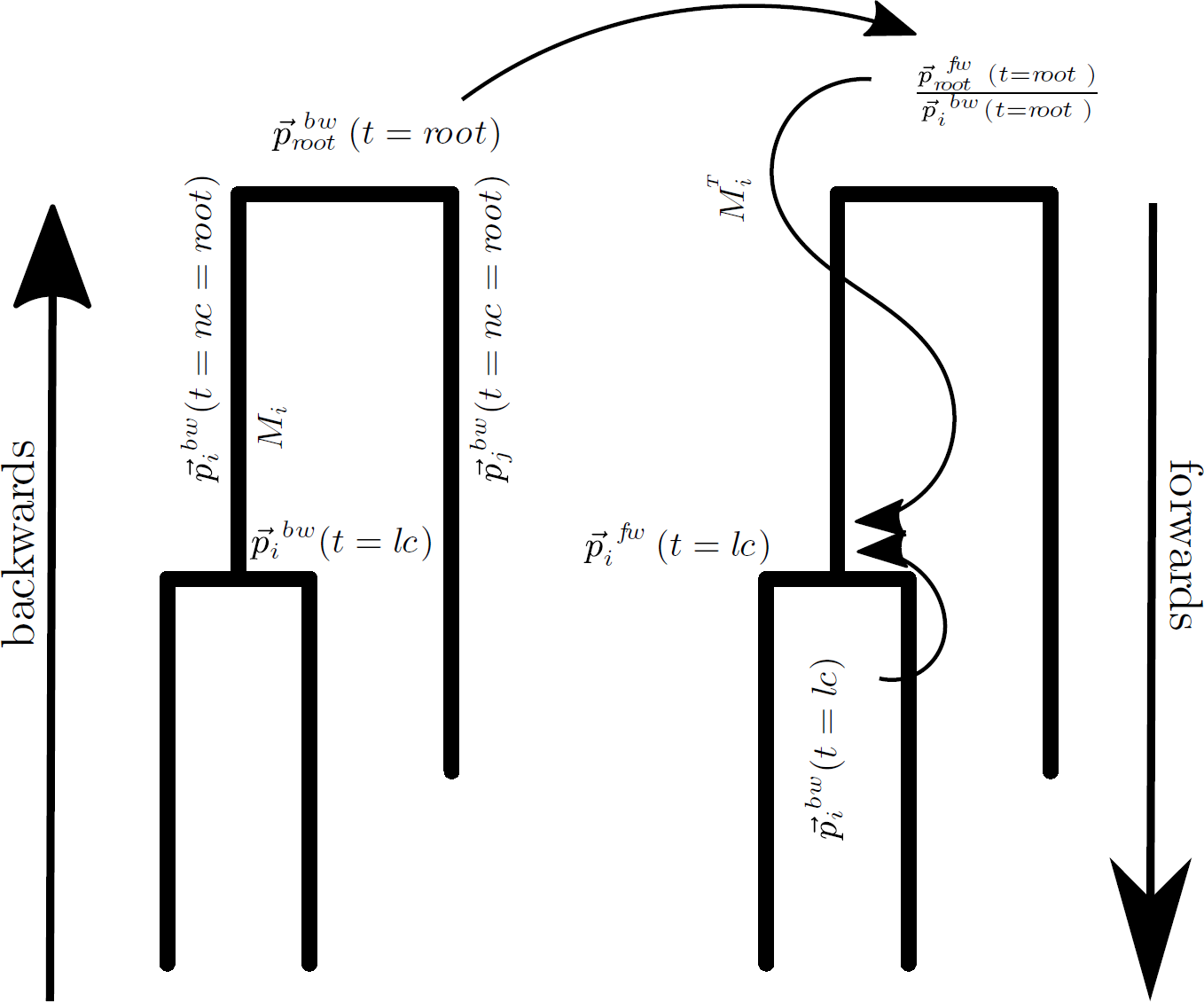
Flow of information using the backwards/forwards algorithm. Going backwards in the tree, we calculate the probability of each node being in any state that includes information up to time *t*. The vector 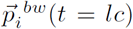 has the entries *P*_*t*=*lc*_(*L*_*i*_ = *a|T*) in position *a*. At the root, the backwards probabilities 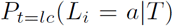 are equal to the forwards probabilities 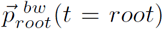. To calculate the downwards probabilities 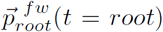, we use the information from all the other parts of the tree and the transition matrix *M*_*i*_ and the backwards probabilities 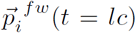 (orange circle)

#### 2.3.3 Forwards calculation of node states including all information in the phylogeny

Based on section 2.2, we know the probabilities of every internal node being in any state. Based on Section 2.3.2, we know how these probabilities change between coalescent events. Going backwards, we calculate 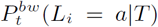, which only includes information up to time *t*. At the root however 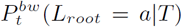 includes information from the full phylogenetic tree from time 0 up to the time of the root. We hence write 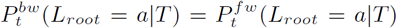, that is the forwards probability of the root being in any state. The forwards probabilities denote the probability of a lineage being in a state that includes information from the full phylogenetic tree. We use the forwards probability 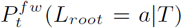 at the root as a starting point to calculate 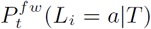 for every lineage *i*. From the root, we proceed forwards in the tree to calculate 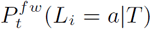 for every internal node at the time *t* of the coalescent event for which lineage *i* was the parent lineage. 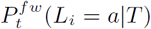 could be calculated at other times as well, we here however focus on the state of nodes. This we do as follows:

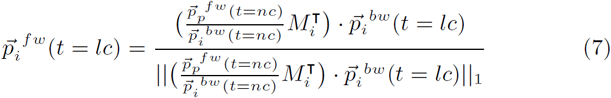

with 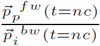 denoting the element-wise division of 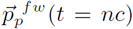, the parent lineage of *i* at the time *nc* of the coalescent event with 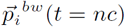, which is the daughter lineage at that time. 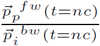 denotes the information of the state of the parent lineage *p* that does not come from lineage *i*. The multiplication with transposed matrix 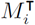 then denotes how much these probabilities have change until the time of the last coalescent event *lc*, where lineage *i* was the parent lineage. The element-wise multiplication with 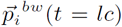 then combines this information with the one from the backwards step. After normalization to ensure that 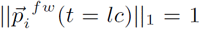, we get the forwards probability of lineage *i* being in any state at time *lc*.

### 2.4 Integration of the differential equations

To integrate equation 1, we use a second order Taylor method with third order step size estimation. This integration technique is similar to the very basic Euler integration, but makes use of the second derivative as well:

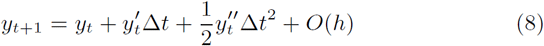

*O*(*h*) stands for derivatives higher than second order. The error that is made by only considering the first and second derivative can be calculated as follows:

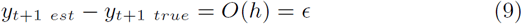

With *y*_*t*+1 *est*_ being the updated term using equation 8 and *y*_*t*+1 *true*_ being the hypothetical true value if all derivatives would be considered. We now assume that the Taylor term of every derivative higher than third are zero. The error *ε* we introduce at every step can therefore be approximated as:

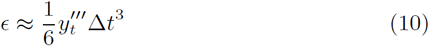

For every integration step, we now choose Δ*t* such that the absolute value of *ε* is smaller than a specified value. While we calculate the second derivative exactly (see supplement), we approximate the third derivative, assuming that the sum of probability mass in each state and that the sum of the derivatives of lineage *i* coalescing in any state is constant (see supplement).

### 2.5 Software

The method above is implemented into our BEAST 2 package MASCOT (Marginal Approximation of the Structured COalsescenT). Simulations were performed using a backwards in time stochastic simulation algorithm of the structured coalescent process using MASTER 5.0.2 (Vaughan and Drummond, 2013) and BEAST 2.4.6 (Bouckaert et al., 2014). Script generation and post-processing were performed in Matlab R2015b. Plotting was done in R 3.2.3 using ggplot2 (Wickham, 2009). Tree plotting and tree height analyses were done using ape 3.4 (Paradis et al., 2004) and phytools 0.5–10 (Revell, 2012). Effective sample sizes for MCMC runs were calculated using coda 0.18–1 (Plummer et al., 2006).

### 2.6 Data availability

The source code of the BEAST 2 package MASCOT is available at https://github.com/nicfel/Mascot.git. All scripts for performing the simulations and analyses presented in this paper are available at https://github.com/nicfel/Mascot-Material.git. Output files from these analyses, which are not on the github folder, are available upon request from the authors. A tutorial is available through the Taming the BEAST project (Barido-Sottani et al., 2017) on how to use MASCOT and its BEAUti interface is available at https://github.com/nicfel/Mascot-Tutorial.git.

## 3 Results

### 3.1 Inference rates and internal node states

First, we want to test how well effective population sizes and migration rates are inferred using MASCOT. We simulated 1000 trees with MASTER (Vaughan and Drummond, 2013) using randomly sampled effective population sizes from Log Normal Distribution(*µ* = −0.125, *σ* = 0.5) and migration rates from an exponential distribution with mean=0.5. We used 1000 tips and 6 different states. The number of tips per state was randomly sampled from a discrete uniform distribution, in order to have scenarios of under- and over-sampling of states. We then inferred the effective population size of every state and the migration rates between each state using MASCOT. The results of these simulations are summarized in figure 2.

**Figure 2:**
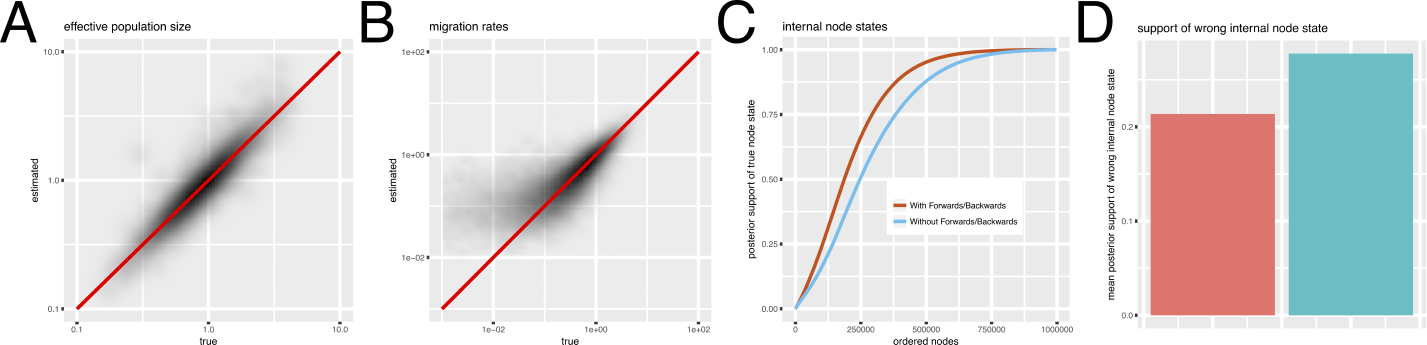
Inference of effective population sizes, migration rates and node states. **A** Inferred migration rates on the y-axis vs. true migration rates on the x-axis. The migration rates between states were sampled from an exponential distribution with mean=0.5. **B** Inferred effective population sizes on the y-axis vs. true effective population sizes on the x-axis. The effective population sizes for the tree simulations were sampled from a lognormal (−0.125,0.5) distribution. The coverage of migration rate estimates was 95% and for effective population size estimates 95.5%. **C** Inferred effective population sizes and migration rates using MASCOT with and without the backwards/forwards algorithm. **D** Mean probability mass allocated to the wrong states of internal node (excluding the root node) with and without the backwards/forwards algorithm

Both effective population sizes and migration rates are inferred well. Population size estimates are however much more precise than estimates of migration rates, see figure 2. This is expected since there are typically much fewer migration events in a phylogeny than coalescent events. Additionally, the number of migration rate parameters estimated (30) is much larger than the number of effective population size parameters (6). The estimates are well correlated with the truth, only at lower migration rates do estimates become worse. This is also to be expected since a low migration rate automatically means less events which will put the estimates closer to the prior (exponential with mean 1). The coverage is 95% for both migration rate estimates and effective population size estimates.

Additional to the inference of parameters, we inferred the state of each internal node with and without the backwards/forwards algorithm. Using the backwards/forwards algorithm reduces the probability mass that is attributed to the wrong node states in this scenario.

### 3.2 Application to H3N2

We then applied MASCOT to 433 Influenza A/H3N2 sequences sampled between 2000 and 2003 from Australia, Hong Kong, New York, New Zealand and Japan. We ran 3 independent chains each for 120 hours using an HKY+ Γ_4_ site model with a fixed clock rate of 5 * 10^−3^ substitutions per site and year. We fixed the clock rate due to a lack of temporal information from the sequences collected for this short amount of time. We then inferred the phylogenetic tree as well as the effective population sizes of every location, the migration rates between them, as well as the additional parameters from the HKY+ Γ_4_ model.

Figure 3 shows the maximum clade credibility tree with the different colors indicating the maximum posterior location estimate of each node. The pie charts indicate the probability of the marked nodes being in any possible location inferred with and without backwards/forwards algorithm. These probabilities are the average over all the node state probabilities for each tree in the posterior containing that clade. We inferred New York to be a source location mainly for strains in Australia and New Zealand. Strains from Japan were inferred to originate mainly from Hong Kong and New York. The root of the phylogeny was inferred to be most likely in New York. The lack of samples near the root however makes the inference of its location unreliable.

**Figure 3:**
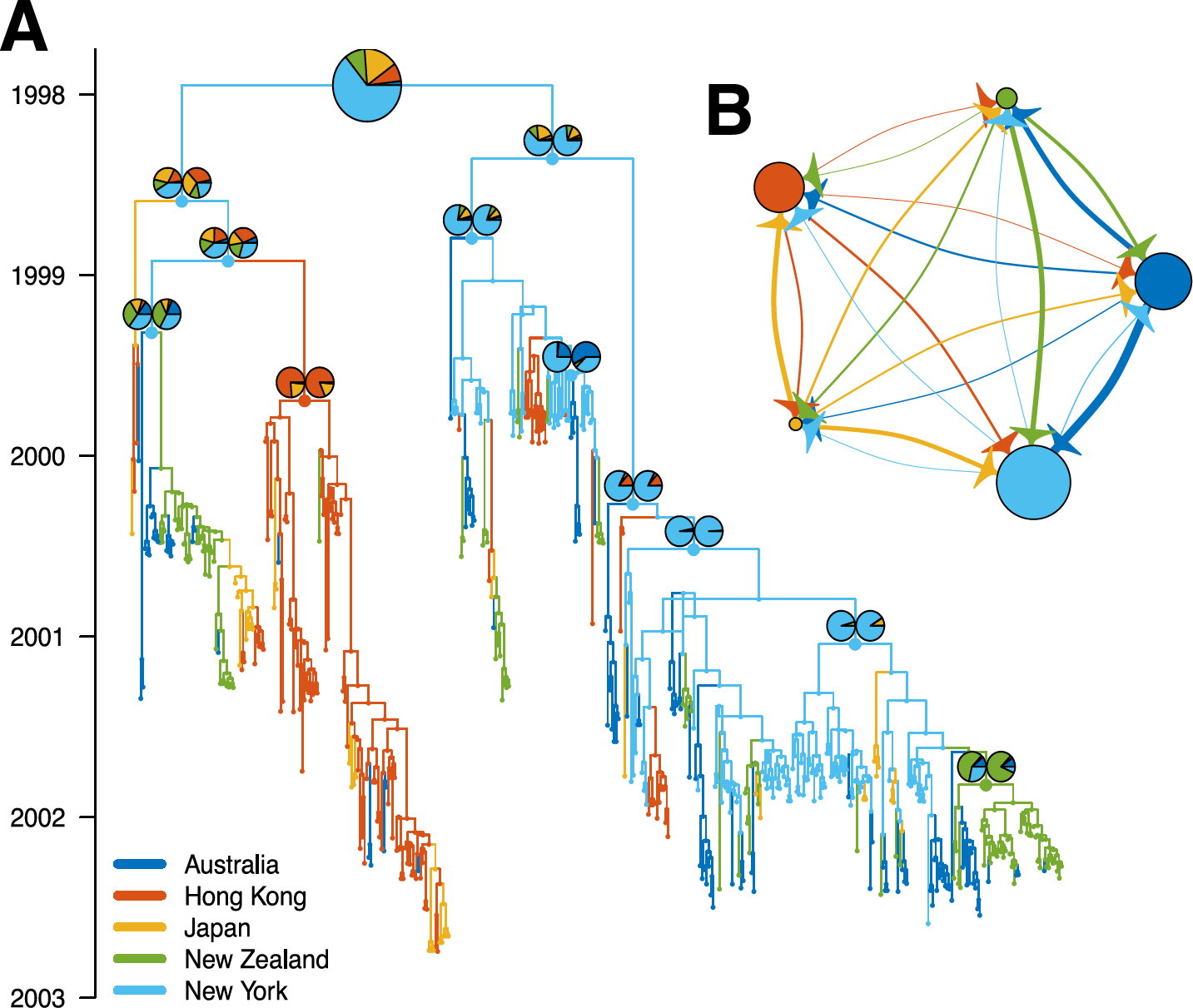
MASCOT analysis of globally sampled Influenza A/H3N2 viruses. **A** Here we show the maximum clade credibility tree inferred from H3N2 sequences from Australia, Hong Kong, New York, New Zealand and Japan. The colour of each branch indicates the most likely state of its daughter node. The pie charts indicate the probability of chosen nodes being in any of the possible states. The left pie chart is the probability inferred using the backwards/forwards algorithm and the right pie chart without using the backwards/forwards algorithm. Since at the root, these probabilities are the same, only one chart is shown. The node heights are the median node heights. **B** The median inferred backwards in time migration rates between locations indicated by the width of the arrow. The different colours of the arrow denote the different source locations, meaning that a green arrow to the red dot shows the backwards in time median migration rate from New Zealand to Hong Kong. The dot sizes are proportional to the median inferred effective population sizes of that state.

## 4 Discussion

We provide a new algorithm to calculate the state of any node in a phylogeny under the marginal approximation of the structured coalescent (Müller et al., 2017). This algorithm entirely avoids the sampling of migration histories. Additionally, we improve the calculation time of our previously introduced approximation to allow for the analysis of phylogenies with more samples and more states.

The calculation time still causes challenges in the analysis of very large datasets. These could be circumvented by a further approximation of 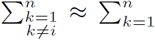 in equation 1 when many lineages are present. This would allow every lineage to have the same transition probabilities and would therefore reduce the number of ODEs that have to be solved.

We then show on simulated data that our approach is able to infer migration rates and effective population sizes reliably even when many different states (6) are present. This is a case where exact methods that sample migration histories are currently not able to reach convergence.

Even though MASCOT is an approximation, we reach a coverage of 95% for migration rates and effective population size estimates. Since this is however still an approximations, there likely are special cases when parameter inference is biased, though we did not find such parameter combinations yet.

We also showed on simulated data that adding an backwards/forwards approach for the calculation of node states improves the inference of internal nodes. We use the backwards/forwards to calculate the state of every internal nodes in a way that is consistent with the complete phylogeny, which is not given by the backwards step alone. In contrast to other approaches (Vaughan et al., 2014; De Maio et al., 2015) we do not explicitly sample nodes states using MCMC. To estimate the probability of a node being in any possible state given a set of parameters we therefore do not need to average over many MCMC samples of migration histories. Whereas for some nodes the difference between with and without forwards/backwards is small, it is especially large for nodes where the difference in where the node is inferred to be compared to the parent node is large.

Finally, we applied MASCOT to a globally sampled H3N2 dataset where we inferred the phylogenetic tree and associated parameters. Our approach is able to reach convergence in the inference of migration rates and effective population sizes, even when a large number of sequences and different locations is present.

MASCOT still requires all migration rates and effective population sizes to be inferred. Especially the number of migration rates (*states* * (*states* − 1)) can become problematic relatively fast. Future additions could however reduce the parameter space by for example deploying Bayesian Search Variable Selection (Lemey et al., 2009) or by making use of generalized linear models (Lemey et al., 2014) to describe migration rates as a combination of different covariates and hence only require the parameters of the GLM model to be inferred.

## Acknowledgements

NM and TS are funded in part by the Swiss National Science foundation (SNF; grant number CR32I3 166258). DR is funded by the ETH Zürich Postdoctoral Fellowship Program and the Marie Curie Actions for People COFUND Program. TS is supported in part by the European Research Council under the Seventh Framework Programme of the European Commission (PhyPD: grant agreement number 335529).

## References

Barido-Sottani, J., Bošková, V., du Plessis, L., Kühnert, D., Magnus, C., Mitov, V., Müller, N. F., Pěcerska, J., Rasmussen, D. A., Zhang, C., et al. (2017). Taming the beast–a community teaching material resource for beast 2–. Systematic Biology.

Beerli, P. and Felsenstein, J. (2001). Maximum likelihood estimation of a migration matrix and effective population sizes in n subpopulations by using a coalescent approach. Proceedings of the National Academy of Sciences of the United States of America, 98(8), 4563–8.

Bouckaert, R., Heled, J., Kühnert, D., Vaughan, T., Wu, C.-H., Xie, D., Suchard, M. A., Rambaut, A., and Drummond, A. J. (2014). BEAST 2: a software platform for Bayesian evolutionary analysis. PLoS computational biology, 10(4), e1003537.

De Maio, N., Wu, C.-H., O'Reilly, K. M., and Wilson, D. (2015). New Routes to Phylogeography: A Bayesian Structured Coalescent Approximation. PLoS genetics, 11(8), e1005421.

Ewing, G., Nicholls, G., and Rodrigo, A. (2004). Using temporally spaced sequences to simultaneously estimate migration rates, mutation rate and population sizes in measurably evolving populations. Genetics, 168(4), 2407–2420.

Hudson, R. R. (1990). Gene genealogies and the coalescent process. Oxford surveys in evolutionary biology, 7(1), 44.

Lemey, P., Rambaut, A., Drummond, A. J., and Suchard, M. a. (2009). Bayesian phylogeography finds its roots. PLoS Computational Biology, 5(9), e1000520.

Lemey, P., Rambaut, A., Bedford, T., Faria, N., Bielejec, F., Baele, G., Russell, C. A., Smith, D. J., Pybus, O. G., Brockmann, D., and Suchard, M. A. (2014). Unifying viral genetics and human transportation data to predict the global transmission dynamics of human influenza H3N2. PLoS pathogens, 10(2), e1003932.

Müller, N. F., Rasmussen, D. A., and Stadler, T. (2017). The structured coalescent and its approximations. Molecular Biology and Evolution, page msx186.

Notohara, M. (1990). The coalescent and the genealogical process in geographically structured population. Journal of Mathematical Biology, 29(1), 59–75.

Paradis, E., Claude, J., and Strimmer, K. (2004). APE: Analyses of Phylogenetics and Evolution in R language. Bioinformatics (Oxford, England), 20(2), 289–90.

Pearl, J. (1982). Reverend bayes on inference engines: A distributed hierarchical approach. In Proceedings of the Second AAAI Conference on Artificial Intelligence, AAAI'82, pages 133–136. AAAI Press.

Plummer, M., Best, N., Cowles, K., and Vines, K. (2006). Coda: convergence diagnosis and output analysis for mcmc. R News, 6(1), 7–11.

Pybus, O. G., Charleston, M. A., Gupta, S., Rambaut, A., Holmes, E. C., and Harvey, P. H. (2001). The epidemic behavior of the hepatitis c virus. Science, 292(5525), 2323–2325.

Revell, L. J. (2012). phytools: an R package for phylogenetic comparative biology (and other things). Methods in Ecology and Evolution, 3(2), 217–223.

Russell, C. A., Jones, T. C., Barr, I. G., Cox, N. J., Garten, R. J., Gregory, V., Gust, I. D., Hampson, A. W., Hay, A. J., Hurt, A. C., de Jong, J. C., Kelso, A., Klimov, A. I., Kageyama, T., Komadina, N., Lapedes, A. S., Lin, Y. P., Mosterin, A., Obuchi, M., Odagiri, T., Osterhaus, A. D. M. E., Rimmelzwaan, G. F., Shaw, M. W., Skepner, E., Stohr, K., Tashiro, M., Fouchier, R. A. M., and Smith, D. J. (2008). The Global Circulation of Seasonal Influenza A (H3N2) Viruses. Science, 320(5874), 340–346.

Stadler, T. and Bonhoeffer, S. (2013). Uncovering epidemiological dynamics in heterogeneous host populations using phylogenetic methods. Philosophical Transactions of the Royal Society B: Biological Sciences, 368(1614), 20120198–20120198.

Takahata, N. (1988). The coalescent in two partially isolated diffusion populations. Genetical research, 52(3), 213–222.

Vaughan, T. G. and Drummond, A. J. (2013). A stochastic simulator of birth-death master equations with application to phylodynamics. Molecular biology and evolution, 30(6), 1480–93.

Vaughan, T. G., Kühnert, D., Popinga, A., Welch, D., and Drummond, A. J. (2014). Efficient Bayesian inference under the structured coalescent. Bioinformatics (Oxford, England), 30(16), 2272–9.

Volz, E. M. (2012). Complex population dynamics and the coalescent under neutrality. Genetics, 190(1), 187–201.

Wickham, H. (2009). ggplot2: Elegant Graphics for Data Analysis. Springer-Verlag New York.

